# A Large-Scale Cryo-EM RNA Motif Dataset and Benchmark for Machine Learning-Based Structure Modeling

**DOI:** 10.64898/2026.02.06.704425

**Authors:** Chandramathi Murugadass, Hajira Rana, Brent M. Znosko, Jie Hou, Dong Si

## Abstract

**Motivation:** RNA molecules play critical roles in gene regulation, viral replication, and cellular control, with their functions tightly coupled to three-dimensional structure. Advances in cryogenic electron microscopy (cryo-EM) now enable RNA structure characterization across a broad resolution range. RNA secondary structural motifs, including hairpins, internal loops, and bulges, act as fundamental building blocks of RNA tertiary architecture and are key targets in RNA-focused therapeutic design. Despite this, most computational approaches for RNA structure prediction from cryo-EM density maps do not explicitly utilize secondary structural motifs as intermediate representations, largely due to the absence of large-scale, high-quality, and motif-resolved datasets suitable for machine learning.

**Results:** Here, we present a large, open-source dataset containing over 125,000 motif-resolved cryo-EM density maps paired with corresponding atomic structures, spanning 25 classes of RNA secondary structural motifs. The dataset covers resolutions from 1.5 Å to 34.0 Å, encompassing both near-atomic and low-resolution density maps relevant to RNA modeling. Each motif instance includes a segmented cryo-EM density map represented as a standardized 3D voxel grid, with atomic-level motif annotations propagated to voxel-level labels for RNA backbone, ribose sugar, and nucleobase components. Segmentation quality is validated via cross-correlation analysis, demonstrating strong agreement between motif-level density maps and atomic reference models. To illustrate the dataset’s utility, high-resolution maps (1.5–2.8 Å) were used to train a machine learning classifier that distinguished five motif classes with a specificity of 0.948.

**Availability and Implementation:** Source code, implementation of the fully automated pipeline, and the benchmark datasets are publicly available at

**GitHub:** https://github.com/DrDongSi/3DEM-RNA-Motif-Dataset

**Zenodo:** https://zenodo.org/communities/3dem-rna-motif-dataset

## Introduction

Ribonucleic acid (RNA) plays essential roles in cellular regulation, viral replication and gene expression, acting both as a functional macromolecule and, in many viruses, as the primary genetic material. Characterizing RNA three-dimensional (3D) conformations and interactions is critical for both biological understanding and therapeutic development (1). Unlike proteins, RNA molecules exhibit substantial conformational flexibility, folding into complex 3D conformations through intramolecular base pairing and long-range interactions. These folds are composed of recurring structural elements, such as helices, hairpin loops, internal loops, bulges, and junctions, that collectively define RNA tertiary structure.

These recurring elements, commonly referred to as *RNA structural motifs*, represent stable and conserved building blocks of RNA architecture (2). By capturing recurring local geometries and interaction patterns across diverse RNA molecules, structural motifs have been widely adopted as fundamental units for RNA structure analysis and modeling. In particular, three-dimensional (3D) structural motifs are commonly used for motif identification, classification, and comparison in curated structural databases such as RNA Characterization of Secondary Structure Motifs (CoSSMos) (3), and serve as motif-level building blocks in RNA structure prediction frameworks based on motif assembly, including VFold (4). Such motif-centric approaches reduce structural complexity while preserving key geometric and interaction features relevant to RNA folding and function.

Recent advances in cryogenic electron microscopy (cryo-EM) have enabled the experimental determination of RNA structures across a broad range of resolutions, providing an increasingly important source of structural data for RNA systems that are challenging to characterize using X-ray crystallography or NMR spectroscopy(5). These cryo-EM density maps offer the opportunity to directly infer RNA structural features from experimental data, motivating the development of computational methods that integrate cryo-EM with structure modeling.

Building on these developments, machine learning and deep learning approaches have shown promise for RNA structure prediction and interpretation from cryo-EM density maps. DeepTracer, ModelAngelo and Cryo2Struct are some popular tools that build atomic-level 3D molecular structures from cryo-EM density maps (6–8). However, progress in this area remains constrained by the limited availability of large, high-resolution experimental training datasets, particularly for full-length RNA structures (9). Full-length RNAs exhibit high conformational variability and structural heterogeneity, complicating consistent modeling and labeling across resolutions and limiting the generalizability of deep learning approaches across RNA families and experimental conditions.

RNA structural motifs provide a natural intermediate representation for cryo-EM density interpretation, bridging atomic models and volumetric data while reducing structural complexity and preserving key local geometric and interaction features essential for RNA folding and hierarchical assembly into tertiary structures; this organization is reflected in Fig. 1, which shows motif-resolved cryo-EM density segments distributed throughout a single RNA structure. By decomposing cryo-EM maps into motif-level density regions, a standardized motif-resolved dataset enables consistent supervision across diverse RNAs and resolutions, supporting motif-aware modeling, supervised learning, and benchmarking directly from cryo-EM data. To address this need, we introduce a large, curated, experimentally derived RNA cryo-EM motif dataset. The dataset comprises 128,018 labeled motif instances spanning 25 RNA secondary structural motif types, with particular emphasis on hairpin loops, bulges, and symmetric and asymmetric internal loops, which represent common and structurally important RNA secondary structure elements. Fig. 2 shows visualization of hairpin structures identified in different RNAs. All motif instances are extracted from experimentally determined cryo-EM maps spanning a wide resolution range. Each motif is paired with its corresponding atomic structure and represented as a standardized 3D voxel grid with voxel-level labels derived from atomic annotations of the RNA backbone, ribose sugar, and nucleobase. This large-scale, motif-resolved cryo-EM dataset with standardized representations and labels enables the development and benchmarking of motif-aware machine learning (ML) methods for RNA structural analysis.

**Fig. 1.**
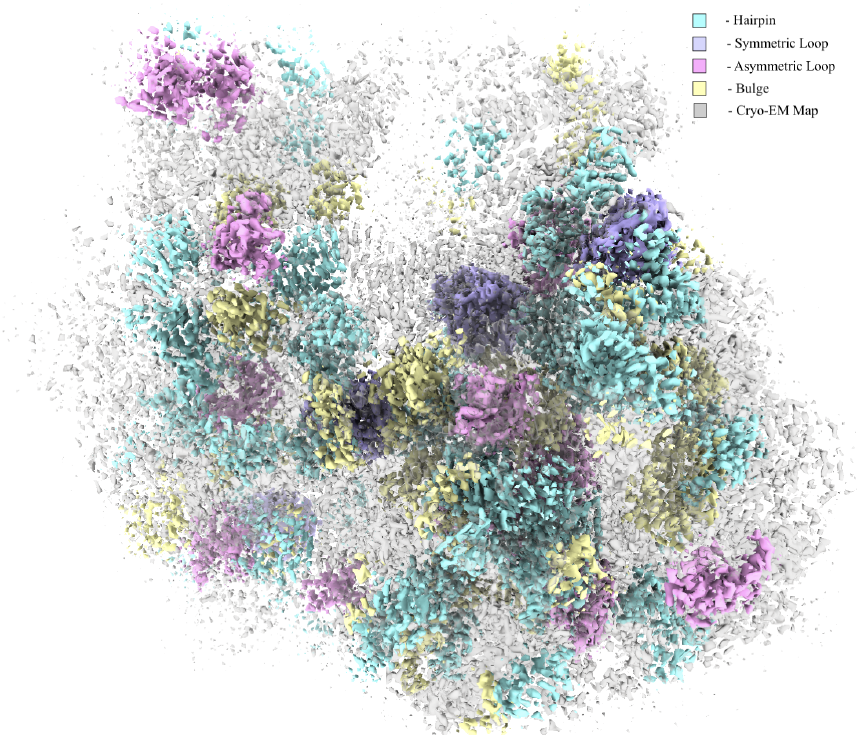
Motif-resolved RNA cryo-EM density maps: Representative motif-level cryo-EM density segments from the *Bacillus subtilis* ribosomal complex (EMDB ID: EMD-18332), colored by RNA secondary structural class. Hairpin loops (cyan), symmetric internal loops (lavender), asymmetric internal loops (pink), and bulges (yellow) are shown. Each segment corresponds to a localized RNA structural motif extracted from the full experimental density map and aligned with atomic-level annotations.

**Fig. 2.**
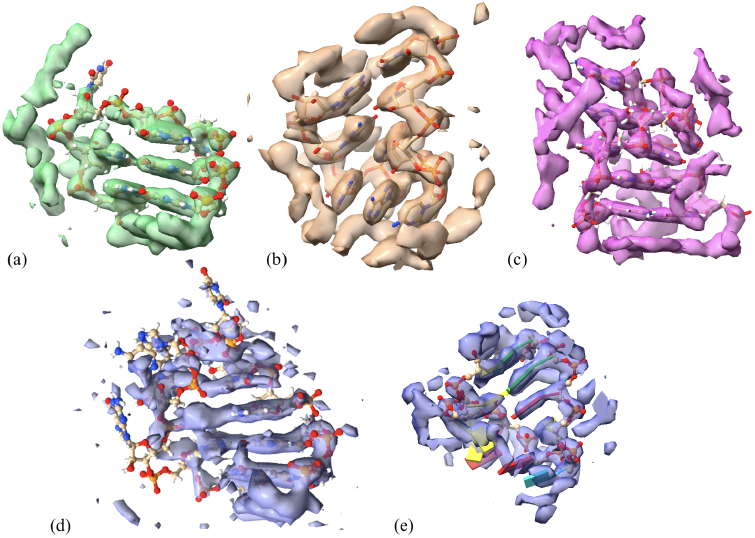
Hairpins: The motif type and its respective PDB ID and global resolution of the source density maps (a) hairpin3 6PJ6 2.20 Å, (b) hairpin4 7PD3 3.40 Å, (c) hairpin5 6PJ6 2.20 Å, (d) hairpin6 6PJ6 2.20 Å, (e) hairpin7 7PD3 3.40 Å

Dataset reliability is ensured through systematic validation of motif-level segmentation using established map–model agreement metrics, including cross-correlation–based fitting scores and atom-level resolvability measures. Additionally, a 3D convolutional neural network for coarse-grained motif classification is developed to demonstrate the dataset’s suitability for supervised learning from cryo-EM density maps. Together, these validations confirm that the dataset provides consistent, high-quality supervision for motif-aware machine learning. At the RNA secondary structure level, the double helix is the core motif, with additional elements classified as hairpin loops as shown in Fig. 2, internal loops (including bulges), and multi-helix junctions. This study specifically targets hairpin loops, bulges, and symmetric and asymmetric internal loops due to their structural and biological relevance.

Symmetric and asymmetric internal loops, as well as bulges, are particularly important as frequent drug-binding sites, making them attractive targets for structure-based drug discovery. Symmetric loops often exhibit more regular geometries, which are advantageous for computational modeling and machine learning–based prediction.

Despite progress, RNA structure prediction remains challenging due to folding complexity, sequence variability, and limited high-resolution experimental structures. As noted by Zhang et al., the scarcity of RNA 3D data has hindered advances compared to protein structure prediction successes such as AlphaFold (9). Existing methods often fail to correctly capture critical motifs, reducing prediction accuracy.

To address these challenges, this work presents a large, curated, experimentally grounded dataset designed to support motif-aware computational methods for cryo-EM based RNA structure prediction. The dataset provides standardized supervision for learning RNA structural features and bench-marking structure prediction approaches.

## Related Works

RNA motif resources and cryo-EM–based structure modeling methods provide important foundations for understanding RNA structure and function. However, existing databases and computational approaches differ substantially in their representation granularity, structural grounding, and suitability for machine learning. Below, we review prior work on RNA motif databases and cryo-EM–based RNA structure prediction, highlighting key limitations that motivate the dataset introduced in this study.

### RNA motif databases

RNA motif databases play a central role in storing recurring sequence and structural patterns that underlie RNA folding and function. Existing resources vary widely in whether they focus on sequence motifs, conserved structural elements, or three-dimensional (3D) motifs derived from experimentally determined structures.

*RNA CoSSMos* provides a curated collection of RNA secondary structure motifs identified from experimentally resolved atomic models (3). The database defines 25 fine-grained motif classes, encompassing hairpin loops of varying lengths, single-nucleotide bulges, and symmetric and asymmetric internal loops with different strand asymmetries. These motifs are systematically classified based on base-pairing patterns and three-dimensional geometry, enabling consistent comparison and analysis across diverse RNA structures. While RNA CoSSMos offers high-quality motif definitions and rich structural metadata, it is derived from atomic coordinates and does not include cryo-EM density maps or volumetric representations. Consequently, RNA CoSSMos is not directly suited for supervised learning from cryo-EM data, but serves as an important reference for defining and categorizing RNA structural motifs. Our dataset complements RNA CoSSMos by extending motif-level annotation into the cryo-EM domain, providing voxel-aligned density representations suitable for machine learning.

*RBPDB (RNA-Binding Protein DataBase)* catalogs RNA–protein binding sequence motifs, typically represented as position weight matrices derived from biochemical experiments (12). While useful for studying RNA–protein interactions, it does not define RNA structural motifs and provides neither atomic coordinates nor cryo-EM density data.

*RNAcentral Expert Databases* aggregate non-coding RNA sequences and annotations from multiple sources (11). Although some entries include conserved secondary structure information, RNAcentral does not explicitly curate RNA structural motifs and does not provide atomic models or cryo-EM density maps suitable for structural or machine learning analyses.

*RNA 3D Motif Atlas (RNA3DHub)* catalogs recurrent RNA 3D structural motifs—such as hairpin loops, internal loops, and junctions—identified from experimentally resolved structures using atomic-level geometric similarity (10). It provides high-quality motif definitions and PDB-derived atomic coordinates but does not include cryo-EM density maps or motif-level volumetric representations.

*MEME Suite* focuses on statistically derived sequence motifs (13), which are not associated with RNA secondary or tertiary structure and lack atomic coordinates or cryo-EM data. Table 1 compares these resources by motif type, structural annotation, atomic coordinate availability, and cryo-EM density. Only RNA CoSSMos and the RNA 3D Motif Atlas provide explicit RNA structural motifs with atomic models. Notably, none of the surveyed databases offer motif-level cryo-EM density maps or voxel-aligned representations suitable for supervised learning, underscoring a key limitation of existing resources for machine learning–based cryo-EM analysis.

**Table 1.**
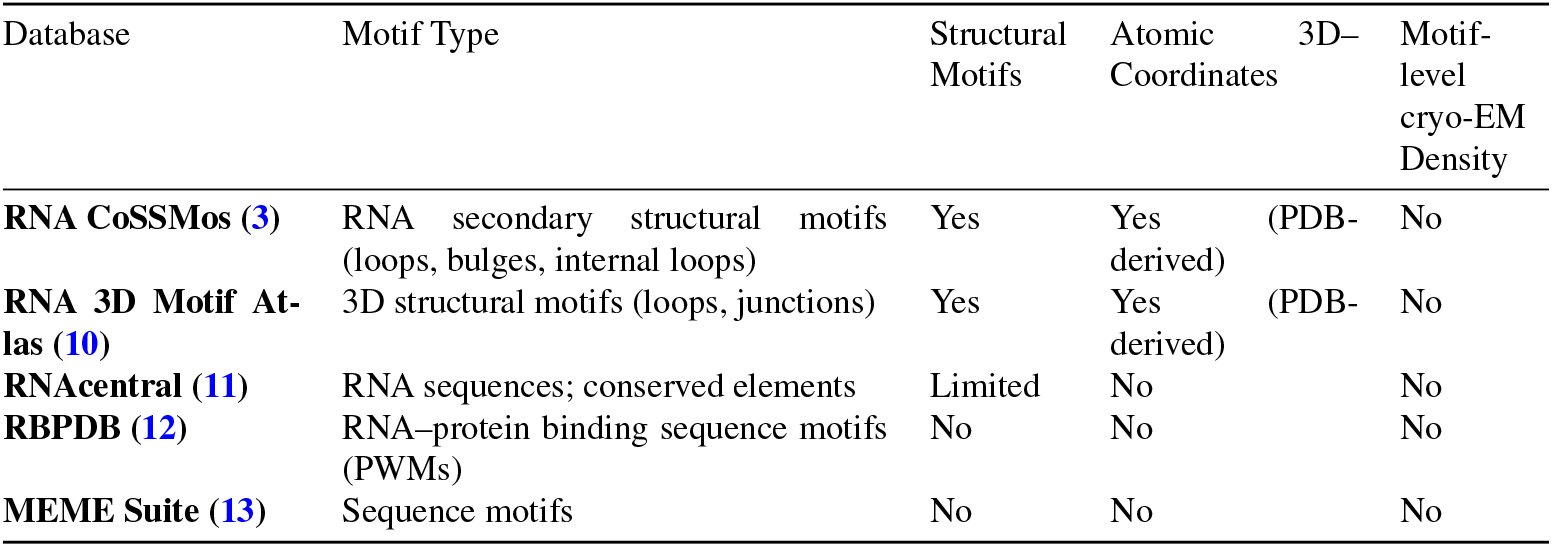
Comparison of RNA motif databases with respect to structural annotation and cryo-EM motif availability.

### RNA structure prediction and modeling from cryo-EM

Machine learning approaches are increasingly applied to cryo-EM density interpretation for RNA, which remains challenging due to conformational flexibility, heterogeneous density, and limited high-resolution data. Most methods therefore rely on careful density preprocessing, accurate alignment between maps and structural labels, and structured postprocessing pipelines.

Sequence-based predictors (e.g., DeepFoldRNA (14), DRFold(15), RhoFold+ (16)) do not directly incorporate experimental density, while physics-based methods such as DRRAFTER rely on heuristic fragment fitting (17). A shared limitation across existing approaches is the lack of large-scale, standardized, motif-resolved cryo-EM datasets to support supervised learning of recurrent RNA structural patterns.

*The dataset introduced in this study is designed to address these limitations by providing motif-resolved cryo-EM density maps with voxel-aligned annotations and systematic quality validation*. By focusing on recurrent RNA secondary structural motifs, this resource enables motif-aware learning and benchmarking directly from experimental density.

## Methods and Materials

### Data Source

The dataset integrates two complementary data types for RNA motif analysis: (1) cryo-EM density maps and (2) RNA secondary structural motif annotations. cryo-EM density maps were obtained from the Electron Microscopy Data Bank (EMDB), which provides experimentally determined volumetric reconstructions across a wide range of resolutions. RNA secondary structure information and corresponding atomic models were sourced from the RNA CoSSMos database and RCSB Protein Data Bank (RCSB PDB), enabling motif-level structural annotation.

As of 2025, the Electron Microscopy Data Bank (EMDB) contains 3,124 cryo-EM maps associated with RNA-containing structures (18). The dataset presented in this study was constructed using the October 2023 release of the RNA CoSSMos database, which provides curated motif annotations for a subset of experimentally resolved RNA structures. Consequently, we curated motif-resolved cryo-EM data from 1,780 RNA cryo-EM maps for which CoSSMos motif annotations and corresponding atomic models were available. Each selected cryo-EM map was paired with its atomic structure to enable motif-level segmentation and labeling. While the current dataset size is constrained by the scope of available CoSSMos annotations, the processing pipeline is fully automated and can be readily re-executed to incorporate future CoSSMos releases and newly deposited RNA cryo-EM structures from EMDB.

Motif extraction relies on three categories of data:

1. RNA motif sequence regions, corresponding to recurring secondary structural elements within RNA chains;
2. Secondary structure metadata, including base-pairing and loop annotations derived from atomic models;
3. cryo-EM density maps, providing experimental volumetric representations of the RNA structures.

Together, these components enable the extraction of motif-resolved cryo-EM density aligned with atomic annotations, forming the basis for downstream analysis and machine learning.

RNA motif definitions, including motif types and residue positions, were obtained from the RNA CoSSMos database, which catalogs three-dimensional RNA secondary structural motifs derived from experimentally resolved atomic models (3).

### Segmentation

Segmentation isolates motif-specific regions from full cryo-EM maps based on atomic structural annotations. For each RNA structure, motif residues were first identified in the atomic model and used to define a spatial neighborhood around the motif. cryo-EM density within a fixed (5.0 Å) radial distance of the motif was extracted to ensure alignment with the selected structural region while minimizing interference from neighboring molecular features.

It begins with loading each RNA atomic structure together with its corresponding cryo-EM density map (Fig. 5). RNA structural motifs are first localized within the atomic model by selecting only the residues that comprise the motif of interest, thereby isolating the relevant structural region while excluding unrelated RNA segments. Once identified at the atomic level, a spatial neighborhood is defined around the motif using a fixed radial distance, capturing the surrounding cryo-EM density associated with the motif. This proximity-based zoning ensures that the extracted density remains structurally aligned with the selected RNA region while minimizing interference from neighboring molecular features.

The zoned density map is then refined by identifying all non-zero density voxels and computing the minimal three-dimensional bounding box that encloses them, effectively removing empty background regions as illustrated in Fig. 3. The resulting cropped volume represents a compact, motif-specific cryo-EM density segment that preserves local structural context while providing a standardized input for down-stream analysis and machine-learning–based RNA structure modeling.

**Fig. 3.**
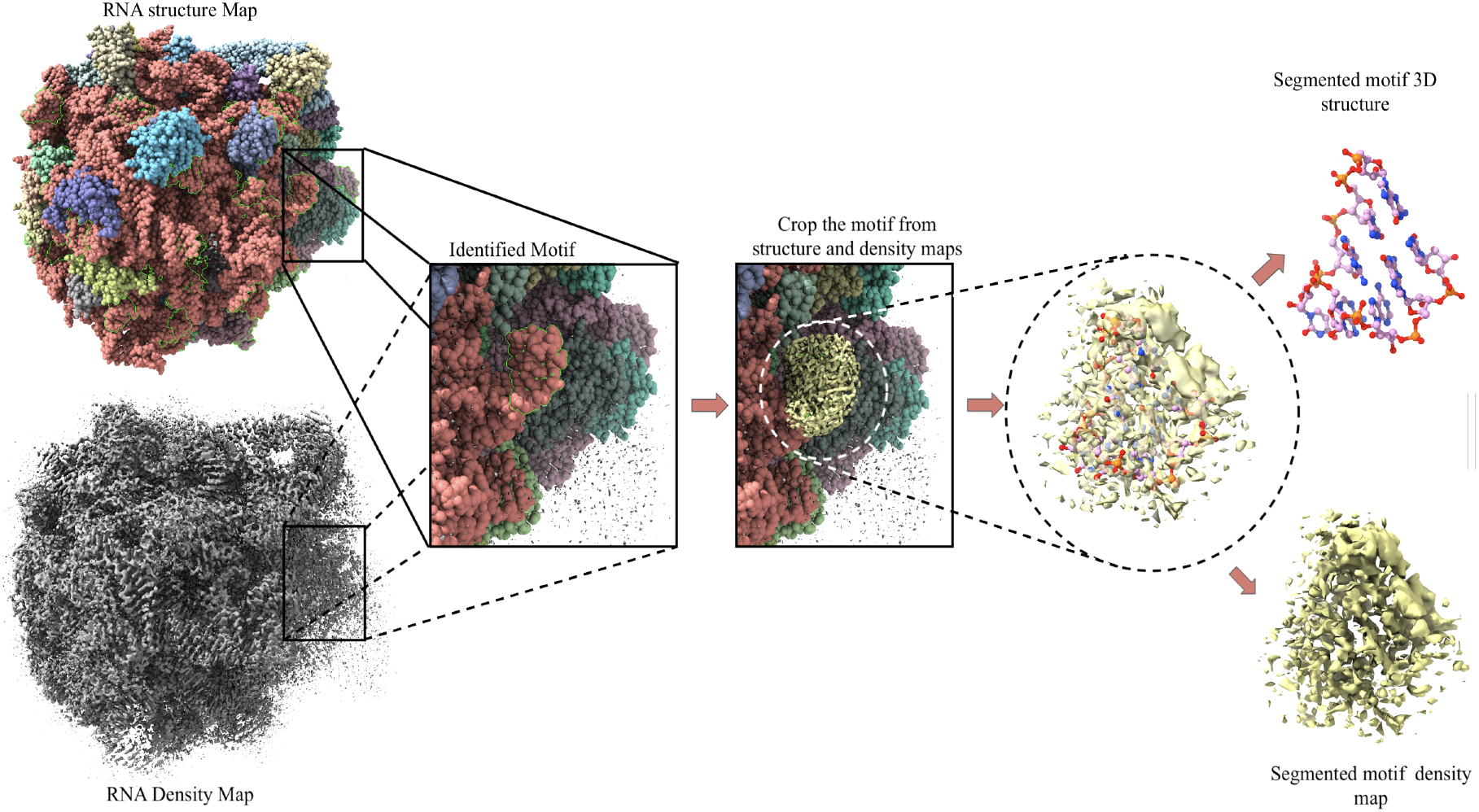
Segmenting a motif from input density map: segmenting a hairpin with four residues at the location 1896 - 1903 chain positions in Chain A of B *subtilis ApdA-stalled* ribosomal complex with PDB ID 8QCQ, by selecting the density voxels around the atomic map of the hairpin.

### Labeling

In the postprocessing stage the density maps are labeled. Prior to labeling the density maps are normalized at a threshold of 95 percentile, we retain only the top 5 percent of the density voxels. In the postprocessing stage, motif-resolved cryo-EM density maps are annotated with voxel-level structural labels derived from corresponding atomic models. Prior to labeling, each density map is normalized using percentile-based thresholding to suppress background noise while retaining high-confidence signal.

Medical Research Council (MRC) density map files are labeled by projecting atomic structural information into the three-dimensional voxel grid of the cryo-EM map. An empty voxel grid matching the dimensions, voxel spacing, and origin of the experimental density map is initialized. Atomic coordinates from the aligned RNA model are then converted into voxel indices, and voxels corresponding to atom centers are labeled according to structural group membership, distinguishing backbone, ribose sugar, and nucleobase components. Since atomic models contain precise coordinate information for residues, secondary structures, or specific motifs, these details can be projected into the volumetric representation of the map. By aligning the atomic model within the density, and initialize an empty voxel grid with the same dimensions as the MRC file and then populate it with labels derived from the model. This provides a structured way to annotate density maps with biologically meaningful features. The labeling process works by systematically traversing the atomic model chain by chain and extracting the coordinates of groups of interest, such as individual atoms, as described in Algorithm 1. These Cartesian coordinates are then converted into voxel indices within the density grid. Then voxels corresponding to the atoms in the respective density maps are marked with a constant value of one. The same steps are performed separately for backbone phosphorus atoms, ribose sugar carbon and oxygen atoms and lastly nucleobase nitrogen, carbon and oxygen atoms creating individual labeled density maps. Fig. 4 shows the three labeled maps of a segmented RNA, where the backbone, ribose sugar, and nucleobase are labeled separately.

**Fig. 4.**
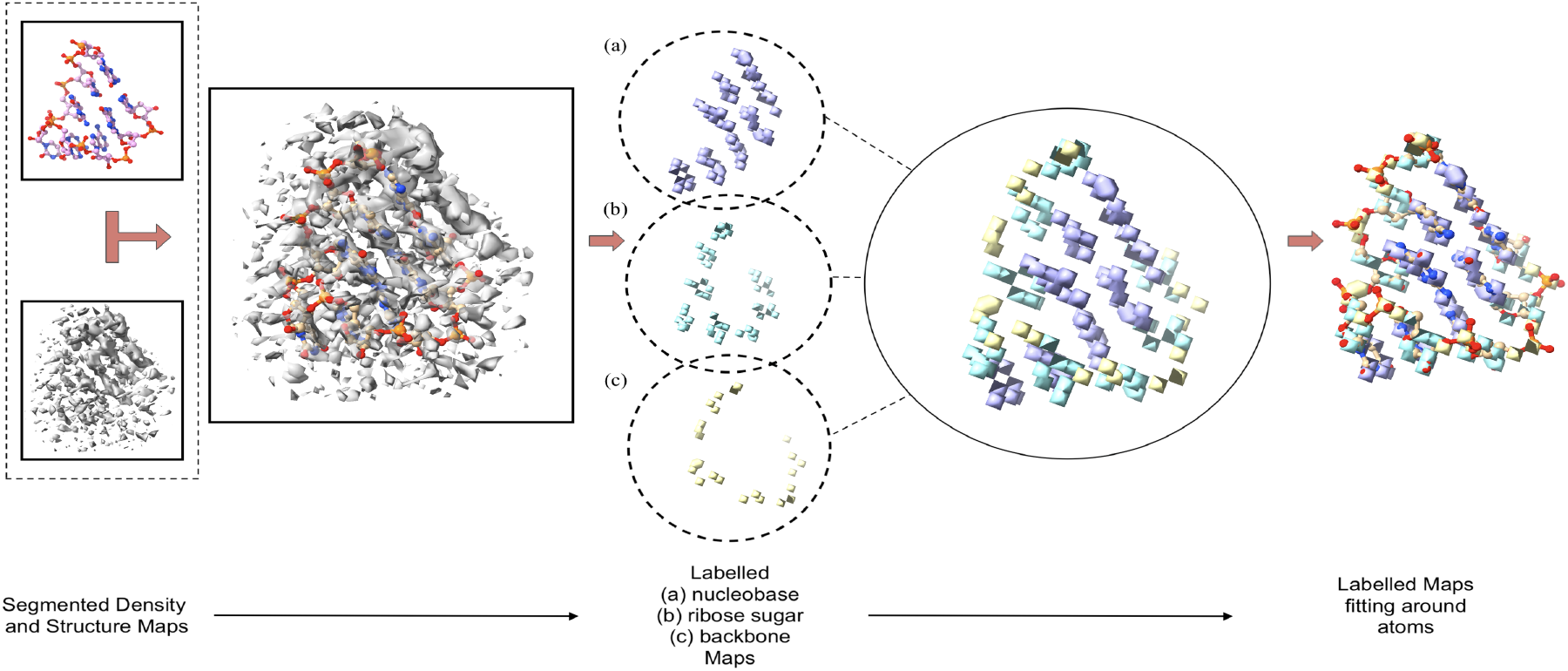
Labelling a segmented motif: Segmented hairpin with four residues at the location 1896 - 1903 chain positions in Chain A of B *subtilis ApdA-stalled* motif is used to generate labels of ribose sugar, nucleobase and the backbone.

**Fig. 5.**
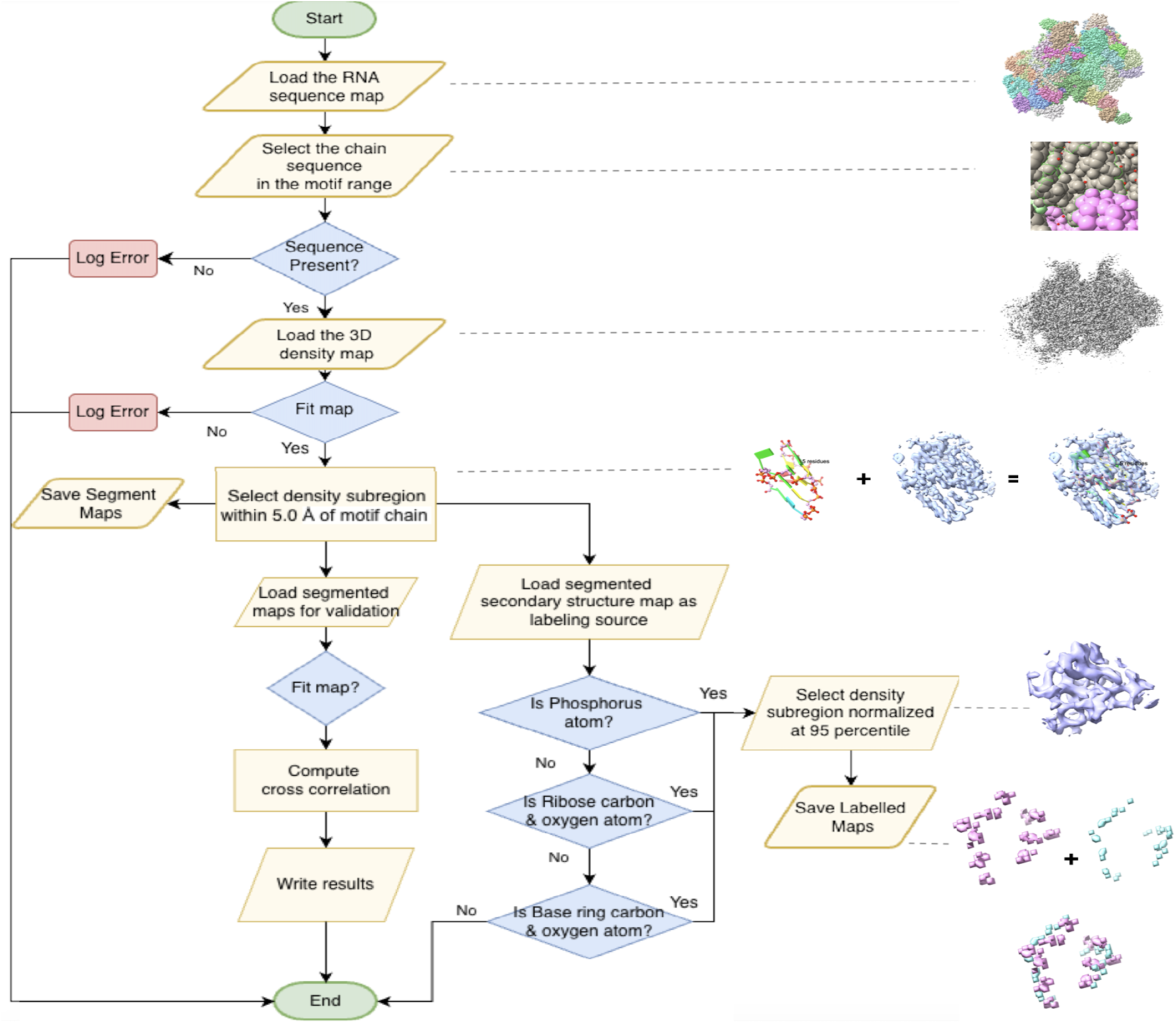
Motif segmentation and labeling: The flowchart depicting the identification and segmentation of motifs from within the larger cryo-EM density maps

### cryo-EM RNA motif classification

To assess the suitability of the dataset for supervised learning, we trained a coarse-grained RNA motif classifier using motif-resolved cryo-EM density maps. Motif annotations were initially defined using the 25 fine-grained RNA secondary structural motif types curated in the RNA CoSSMos database, encompassing hairpin loops, bulges, and symmetric and asymmetric internal loops of varying sizes.

For classification, these 25 fine-grained motif types were deterministically grouped into five biologically meaningful classes to reduce label sparsity and improve learning stability: *symmetric internal loops, asymmetric internal loops, hairpins, bulges*, and *unknown*. Symmetric loops include canonical *n*×*n* internal loops with *n* = 1 … 5, while asymmetric loops comprise non-equal *m*×*n* internal loop pairs with 1≤*m, n*≤5 and *m* ≠ *n*. Hairpins were defined by loop lengths of 3–7 nucleotides, and bulges correspond to single-sided insertions of length 1–5 nucleotides. Each motif instance was mapped from its original RNA CoSSMos annotation to one of these grouped classes. We also consider ‘Unknown’ group which refers to density maps comprising non-motif segments extracted from the cryo-EM densities that act as background or noise.

Density maps were resampled to a uniform voxel size, normalized, and cropped into fixed-size 64^3^ voxel patches centered on motif regions. Classification was performed using a 3D convolutional neural network comprising three convolution–ReLU–max pooling blocks as shown in Fig.6, followed by two fully connected layers. The network was trained using cross-entropy loss and the Adam optimizer, with five-fold cross-validation used for performance assessment (Table 2). Model architecture and training details are summarized in Algorithm 2.

**Fig. 6.**
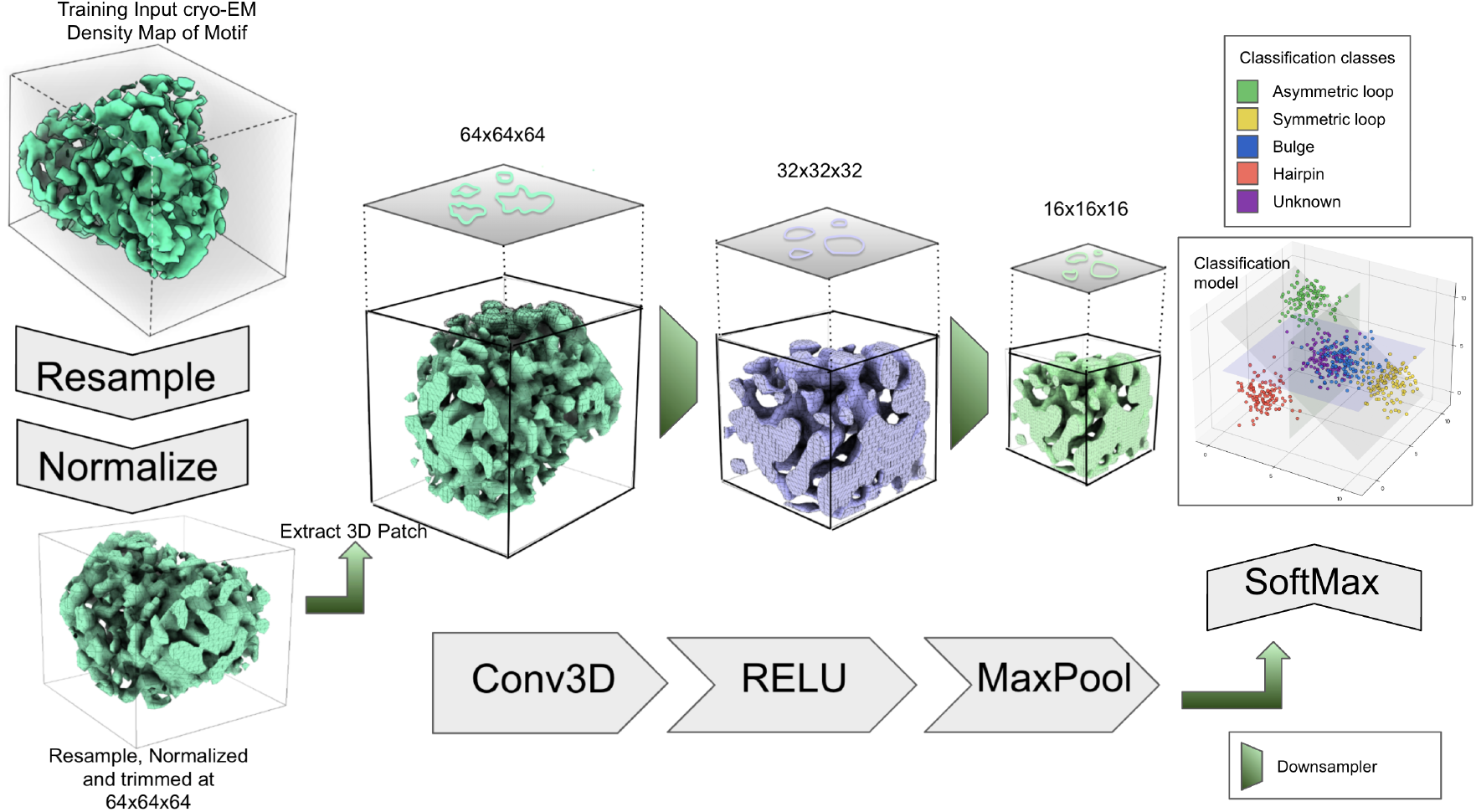
3D CNN Classifier Workflow: Takes segmented density maps as input and builds a classification model that classifies the known motif data into five motif classes.

**Table 2.**
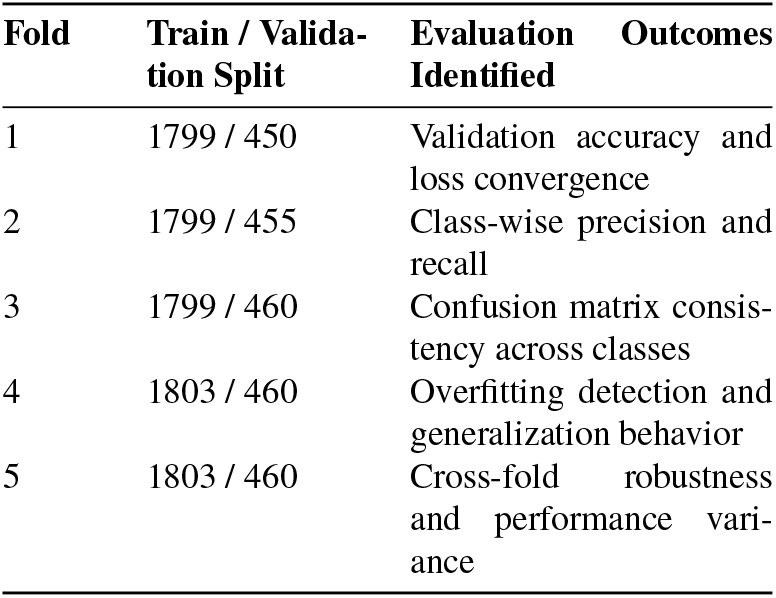
Five-fold cross-validation scheme used for training and evaluating the RNA motif classifier. All folds were evaluated using accuracy, class-wise precision and recall, confusion matrices, and specificity.

#### Algorithm 1 RNA cryo-EM Density Labeling from Atomic Structures

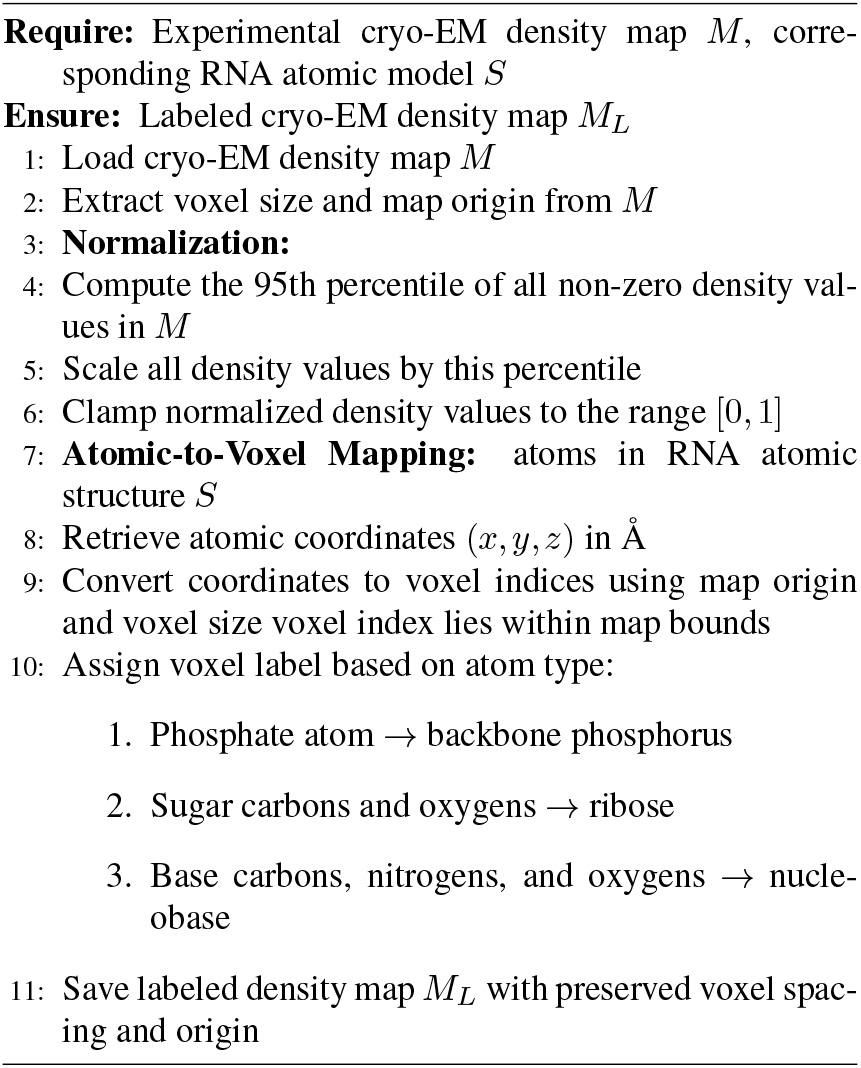

cryo-EM density maps were resampled to a uniform voxel size, normalized, and cropped into fixed-size 64 ×64× 64 voxel patches centered on motif regions. These patches served as input to a 3D convolutional neural network (CNN) consisting of three convolution–ReLU–max pooling blocks followed by two fully connected layers. The network outputs class probabilities over the five grouped motif categories and was trained end-to-end using cross-entropy loss and the Adam optimizer.

Model performance was evaluated using five-fold cross-validation, with equal-sized splits used to assess classification accuracy, robustness across folds, and class-wise performance consistency (Table 2). Architectural and training details are summarized in Algorithm 2. The objective of the model is to learn discriminative three-dimensional density patterns corresponding to coarse-grained RNA structural motif classes from cryo-EM volumes. Each input density map is first resampled to a uniform target voxel size and normalized to zero mean and unit variance. From the resampled volume, a fixed-size cubic patch of dimensions 64×64×64 is extracted and represented as a single-channel tensor *D* ∈ℝ^1×64×64×64^. Fine-grained RNA motif annotations spanning 25 motif types are deterministically grouped into five classes: symmetric loops, asymmetric loops, hairpins, bulges, and unknown motifs.

The density patch is processed using a three-layer 3D convolutional neural network. The feature extractor consists of three convolutional blocks, each composed of a 3×3×3 convolution, a ReLU nonlinearity, and 2×2×2 max pooling as shown in Fig. 6. The channel dimensions progress as 1→ 16→ 32→ 64, progressively capturing higher-level spatial features while reducing spatial resolution. The resulting feature map is flattened and passed through a fully connected classifier with a 128-unit hidden layer and a final linear layer producing logits over five motif classes.

The network is trained end-to-end using the Adam optimizer with a learning rate of 10^−3^ and cross-entropy loss. Minibatches of size four are used, and validation accuracy is monitored after each epoch to assess generalization performance.

#### Algorithm 2 Training 3D CNN for RNA Motif Group Classification

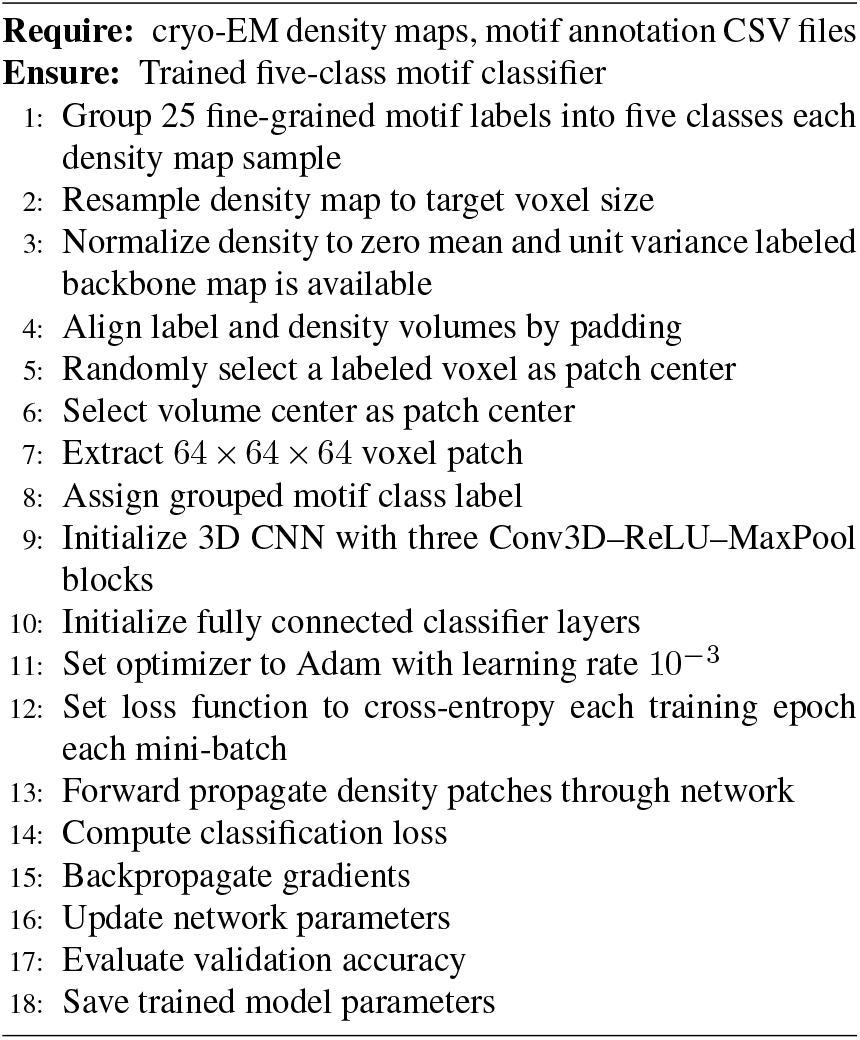

### Validation

The accuracy of motif-level segmentation and labeling was evaluated using established model–map fitting metrics that quantify agreement between atomic structures and cryo-EM density. Two complementary measures were employed, the real-space cross-correlation and atom-level Q-scores. Guidelines can be included for standard research article sections, such as this one.

#### Metrics

##### 1. Cross Correlation

In cryo-EM, a Cross Correlation is a metric that quantifies how well an atomic model fits a density map, indicating the likelihood of a correct fit for a given feature. It measures the difference between the cross-correlation score of the feature at its placed location versus its score at displaced positions. The scores are in the range of [0,1], with a higher score meaning a better, more resolved fit (19). A quantitative measure to what degree structural features are actually resolved in cryo-EM density maps.

The atomic model to the density map fit is typically evaluated according to the Cross Correlation of two maps constructed from independent data sets. The research used Phenix software that provides a command phenix.map_model_cc to perform map to model correlation. The model–map correlation coefficient (model_map_CC) is a real-space metric that quantifies how well an atomic model agrees with an experimental density map. The model-map correlation coefficient (model_map_CC) is a real-space metric used to quantify the agreement between an atomic model and an experimental density map. In Phenix, the correlation is computed between the experimental map and a model-derived map sampled on the same grid, with the model density calculated from atomic coordinates, occupancies, atom types and displacement parameters. For cryo-EM maps, which frequently have non-zero mean density values, Phenix employs a mean-subtracted, normalized correlation formulation that measures covariation rather than absolute density overlap. The correlation coefficient is defined as

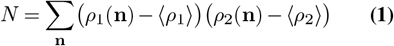

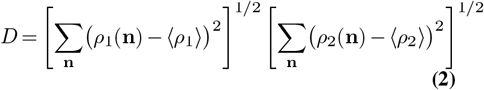

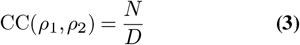

where *ρ*_1_ and *ρ*_2_ denote the experimental and model-derived maps, respectively, and the summation is performed over selected grid points **n**. The interpretation of CC depends critically on the region used for its calculation; CC_mask_ restricts the evaluation to voxels within a molecular mask surrounding the model, thereby excluding solvent and background noise and providing a more localized and physically meaningful assessment of model fit. CC_mask_ is therefore commonly reported as a robust indicator of model accuracy in cryo-EM structure validation.

##### 2. Q-score

The Q-score calculates the resolvability of atoms by measuring similarity of the map values around each atom relative to a Gaussian-like function for a well resolved atom. Q-score of 1 indicates that the similarity is perfect where a small value of Q-score which is closer to 0 indicates the similarity is low (20). The research used ChimeraX command line tool extension that supports Q-score computation. It does this by defining a series of spherical shells around the atom at intervals of shellRadiusStep up to maxShellRadius, and attempting to find pointsPerShell points in each shell that are closer to the test atom than any other atom in the model (21).

The command also comes with an option to write the Q-score computation metadata to the file hence the computed score is present as a labeled metadata on the file.

Model-to-map fit can also be evaluated in Fourier space by calculating the correlation between Fourier map coefficients binned in resolution shells. The calculated CC values are typically represented as a function of the inverse of resolution and are called the Fourier shell correlation (FSC) (19).

#### Cross-correlation

Model–map agreement was assessed using the real-space cross-correlation coefficient (CC), which quantifies the similarity between an experimental cryo-EM density map and a model-derived map sampled on the same voxel grid (20, 22). CC values range from 0 to 1, with higher values indicating better structural agreement and resolvability.

Cross-correlation was computed using the Phenix tool `phenix.map_model_cc` (19), which applies a mean-subtracted, normalized formulation suitable for cryo-EM maps with non-zero background density. To obtain localized and physically meaningful assessments, correlations were calculated within a molecular mask surrounding each motif (CC_mask_), excluding solvent and background regions. CC_mask_ is widely reported as a robust indicator of model accuracy in cryo-EM structure validation (23).

#### Q-score

Atomic resolvability was further evaluated using Q-scores, which quantify the agreement between the atomic model and the experimentally derived cryo-EM density map. Q-scores assess how well each atom is supported by the local density by comparing the atom-centered density profile to an idealized Gaussian reference (21). Q-scores range from -1 to 1, with higher values indicating better atomic-level resolution and consistency with the density map.

#### Fourier shell correlation

Model–map agreement was evaluated using Fourier shell correlation (FSC), which measures how well a density map calculated from an atomic model matches the experimental cryo-EM map across different spatial frequencies (19). The two maps are compared in Fourier space by grouping their coefficients into resolution shells (concentric bands corresponding to narrow ranges of spatial resolution) and computing the correlation within each shell. FSC analysis was performed on representative motif instances to complement CC_mask_ and Q-score–based local validation. FSC values are plotted as a function of inverse resolution, with higher values indicating stronger model support by the experimental density and declines at higher frequencies reflecting resolution limits and reduced agreement at finer structural detail.

The Fig.7 shows the FSC correlation between model and map. The model–map FSC demonstrates strong agreement between the atomic model and the experimental cryo-EM density across low and intermediate spatial frequencies, with masked FSC values exceeding unmasked values, indicating a well-supported model within the molecular region and map-limited resolution at high spatial frequencies.

**Fig. 7.**
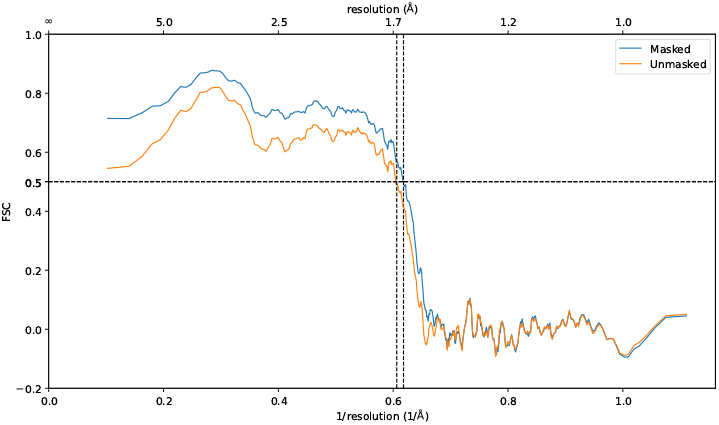
Model-map fourier shell correlation (FSC): Masked and unmasked FSC curves comparing the atomic model with the experimental cryo-EM density map as a function of spatial frequency. The masked FSC is computed over the molecular region, whereas the unmasked FSC is computed over the entire map volume. Results are shown for the *Escherichia coli* ribosome (PDB ID: 8B0X), focusing on a representative 1×3 internal loop motif from chain A (nucleotides 1320–U, 1321–C, 1322–C, 1323–A, 1324–A, 1325–C, 1326–G and 1332–C, 1333–G, 1334–G, 1335–G, 1336–G).

**Fig. 8.**
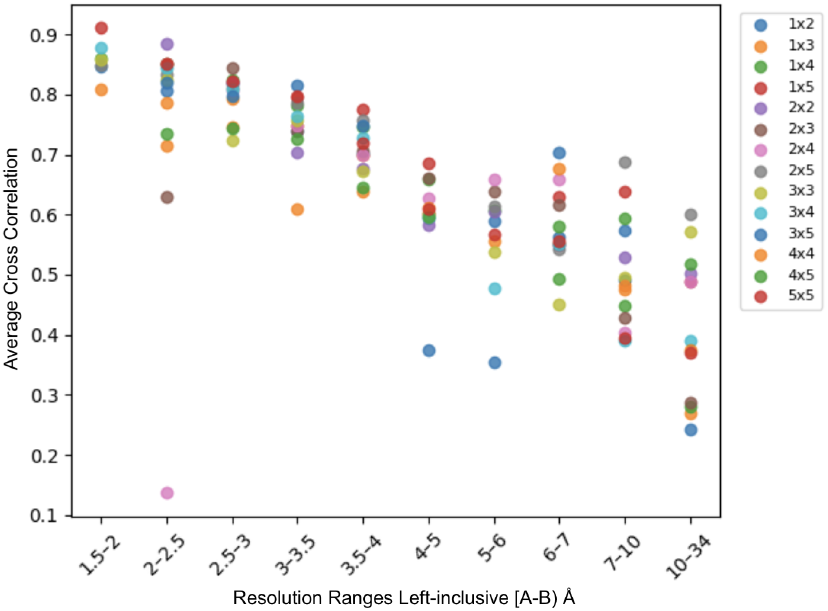
Average cross correlation: CC_mask_ for internal loops at varying resolutions.

**Fig. 9.**
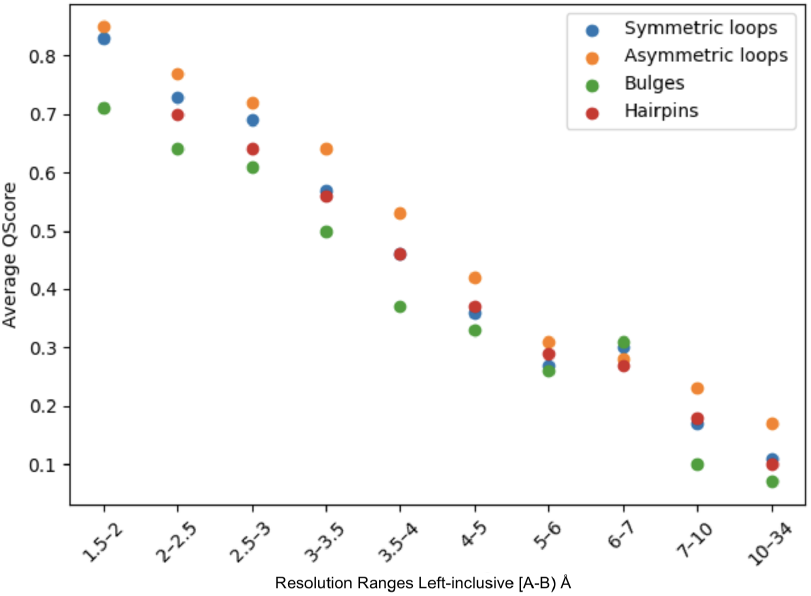
Average Q-scores: map-to-model fitting of motifs of type hairpin, bulges, symmetric and asymmetric loops

## Results

Fitting score measures how well a segmented RNA structure agrees with an experimentally derived cryo-EM density map. By calculating the correlation between the 3D motif placements and the actual electron density, we can validate whether each RNA segment is correctly positioned. A high fitting score (typically *>*0.8) indicates accurate spatial alignment, making it a valuable metric for assessing motif assembly quality and structural plausibility.

**Atom/Model-to-map agreement** was evaluated using the Q-score metric, which quantifies the consistency between the generated atomic model and the corresponding cryo-EM density map. The metric was evaluated across more than 12,500 randomly selected samples. By assessing local density patterns around individual atoms, Q-scores provide a stable measure of atomic resolvability that is robust to global intensity scaling and map normalization. As such, Q-scores offer a reliable indicator of how well atomic coordinates are supported by the experimental density. Analysis of the correlation trends in Tables 3 and 4 reveals that certain internal loop motifs exhibit enhanced robustness as resolution decreases. Notably, the 2 × 5 motif maintains relatively high correlation values, remaining near 0.78 at 3.0 Å and retaining approximately 0.61 even at coarser resolutions of 7–10 Å. Likewise, the 3 × 3 motif consistently outperforms other 3× N configurations across resolutions. This behavior likely reflects the geometric regularity of these motifs and the presence of distinctive, well-preserved density features that remain detectable under reduced resolution. Figs. 8 and 9 show the pattern of cross correlation and Q-score values for different motif types at varying resolutions of the density map.

**Table 3.** Cross correlation trends across cryo-EM resolution ranges.

| Resolution (Å) | CC Range | Observed Pattern |
| --- | --- | --- |
| 1.5–2.5 | 0.80–0.90 | Consistently high correlation |
| 3.0–4.0 | 0.70–0.75 | Moderate but stable correlation |
| 5.0–10.0 | 0.50–0.60 | Noticeable degradation |
| 10.0–34.0 | 0.24–0.40 | Significant loss of correlation |

**Table 4.**
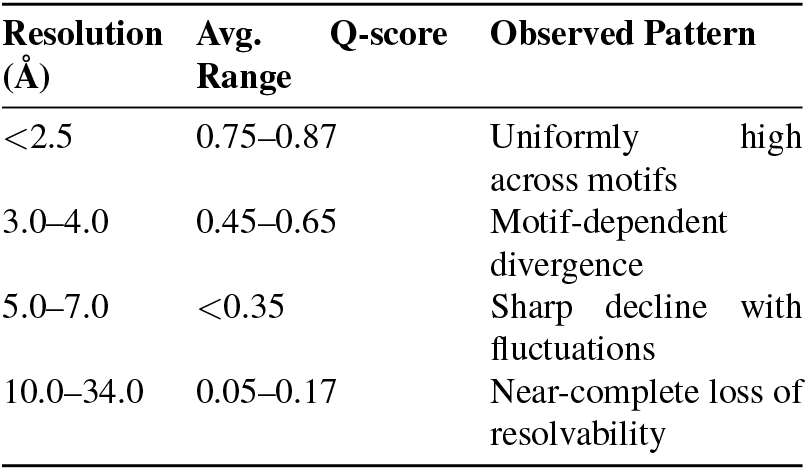
Average Q-score correlation trends across cryo-EM resolution ranges.

**Table 5.**
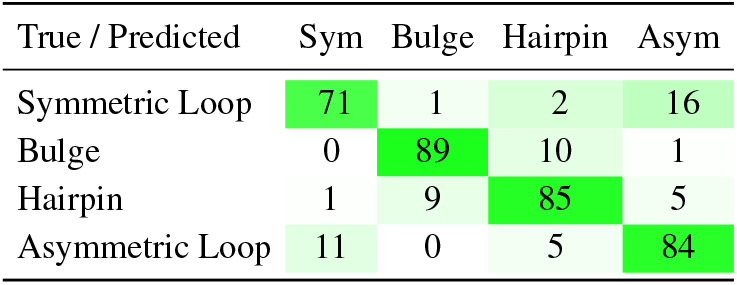
Combined confusion matrix across all motif evaluations. Rows indicate true labels and columns indicate predicted labels.

**Table 6.** Per-class sensitivity and specificity of the 3D CNN motif classifier, computed from the combined confusion matrix.

| Motif Class | Sensitivity | Specificity |
| --- | --- | --- |
| Symmetric loop | 0.789 | 0.960 |
| Asymmetric loop | 0.840 | 0.924 |
| Hairpin | 0.850 | 0.941 |
| Bulge | 0.890 | 0.966 |
| **Macro average** | **0.842** | **0.948** |

As shown in Fig. 7, masked FSC values exceed unmasked values across low and intermediate frequencies, indicating strong support for the fitted atomic models within the molecular regions and resolution-limited agreement at higher fre-quencies.

The trained 3D CNN was evaluated on randomly sampled motif patches using ground truth labels and a test set containing more than 350 samples. Performance was assessed separately for symmetric loops, asymmetric loops, hairpins, and bulges using confusion matrices, sensitivity, and specificity.

For symmetric loops, the model achieved a sensitivity of 0.789, indicating strong detection capability, though misclassifications primarily occurred as asymmetric loops. Asymmetric loop detection showed a sensitivity of 0.840, with most errors arising from confusion with symmetric loops and asymmetric loops. Hairpin motifs were classified with a sensitivity of 0.850, demonstrating robust recognition despite occasional mislabeling as bulges or asymmetric loops. Bulge motifs exhibited a sensitivity of 0.890, reflecting reliable detection with limited confusion.

Across all evaluations, non-target class specificities were generally high (0.90–1.00), indicating effective rejection of incorrect classes. Overall, the results suggest that the model learns discriminative structural features for each motif type, while highlighting the need for balanced multi-class evaluation to improve global performance.

### Inferences

In conclusion, based on the cross correlation results it can be inferred that:

1. Fitting score is highest for high resolution maps on average greater than 0.85 score for both Q-score and CC_mask_ correlation which is a very high measure of fitness.
2. For symmetric motifs, with smaller motifs (1×1, 2×2, 3×3) maintain better fit with decreasing resolution, while 4×4 deteriorates more quickly with decreasing resolution.
3. Simple motifs (1xN) are consistently the easiest to detect and maintain higher scores across resolutions.
4. Larger motifs, such as bulges of five nucleotides, have much lower average correlations overall even at high resolutions compared to the overall correlations of smaller motifs at the same resolution.
5. At very low resolutions (10–34 Å), all motifs perform poorly, indicating insufficient structural detail.

## Discussions

More than 98% of the metadata collected for segmentation was successfully segmented except for a few outliers, some of the outliers that were dropped from segmentation were because of improper fitting in the RNA map to model. For example, as shown in Fig. 10 (a) has the atomic model poorly fitting the cryo-EM map, such outliers were dropped from segmentation and hence absent in the final dataset.

**Fig. 10.**
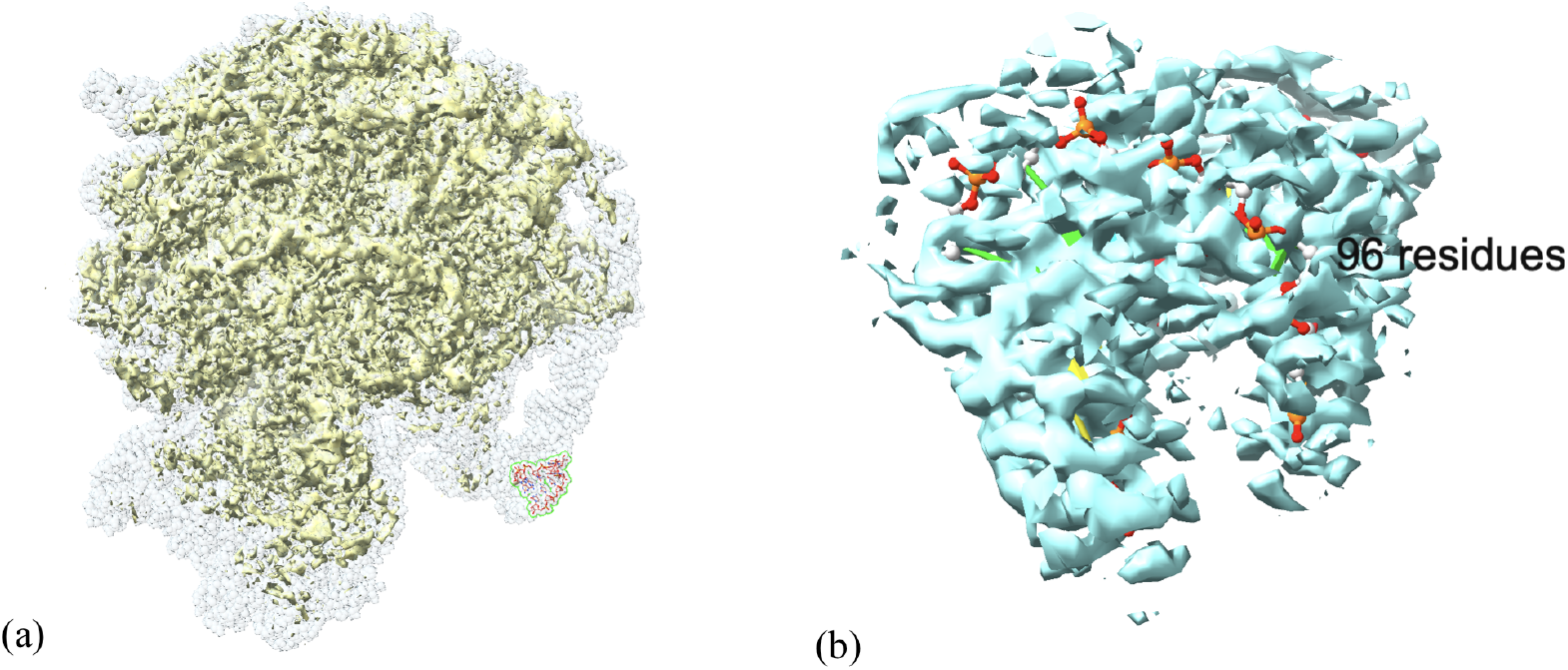
Outlier motifs: (a) 70S ribosome from *Mycobacterium tuberculosis* bound with capreomycin with PDB ID 5V93 and EMD ID EMD-8645 having the motif segment outside the available density map (b) Segmented 2×3 motif of erythromycin resistant *Staphylococcus aureus* 70S ribosome at 234-A 235-G 236-A 237-U 238-G 239-G 131-C 132-C 133-U 134-A 135-C 136-C 137-U

Similarly, it can be seen that motif type 2×3 segmentation has unusually low cross correlation value at 2.0–2.5 Å 0.629 but high at 2.5–3.0 Å (0.843), this is attributed to the fact that some segmented motif of RNAs such as erythromycin resistant *Staphylococcus aureus* 70S ribosome have low fitting score cross correlation value of 0.2617 and Q-score value of 0.15. As seen in Fig. 10 (b) the atomic model can be seen visibly outside most density voxels, and also the Q-score of the original ribosome to the density map is itself a low fitting score of 0.49. Source map to model fitness was poor hence the fitness of the segmented sample is also poor.

In addition to the segmentation validation, we tested the models trained over each fold shown in Table.2 and found the following pattern. Fold 1 demonstrated stable optimization behavior, with training loss steadily decreasing and validation accuracy converging to 0.86, indicating effective learning without instability. Fold 2 showed strong and balanced class-wise performance, achieving high macro sensitivity (0.85) and specificity (0.96), with particularly robust detection of bulge motifs. Fold 3 demonstrated consistent class discrimination across all motif categories, achieving a macro sensitivity of 0.84 and macro specificity of 0.96, with uniformly high per-class performance and minimal confusion among structurally distinct motifs. Fold 4 revealed minimal overfitting, with validation accuracy stabilizing around 0.81 despite continued loss reduction, suggesting good generalization behavior under an alternate data split. Fold 5 confirmed cross-fold robustness, achieving the highest macro sensitivity (0.86) and specificity (0.96), demonstrating low performance variance and consistent motif classification across splits. The classification models validation accuracy showed a consistent upward trend across all folds, while the training loss rapidly decreased and converges to near-zero values, indicating stable optimization and effective convergence of the model during training. Although the training loss became extremely small in later epochs, the validation accuracy plateaus rather than declining, suggesting minimal overfitting and demonstrating that the model maintains good generalization to unseen validation data. Across the five cross-validation folds, the final validation accuracies are tightly clustered in the range of approximately 0.82 to 0.86, reflecting strong cross-fold robustness and low performance variance, which indicates reliable and stable model behavior.

In conclusion, this study demonstrates that RNA 3D structural motifs can be reliably extracted from cryo-EM density maps with consistently acceptable model–map agreement. We further show that these motif-resolved density representations can be effectively leveraged by machine learning algorithms, establishing their suitability for supervised learning from experimental cryo-EM data. The openly released dataset provides a broadly applicable foundation for the development, evaluation, and benchmarking of RNA-focused machine learning methods, and is expected to facilitate continued advances in data-driven RNA structural analysis.

## Author contributions statement

J. Hou and D. Si conceived the idea of this work. C. Murugadass conducted the research, curated the data, and developed the code. C. Murugadass and D. Si developed the method and analyzed the results. C. Murugadass and H. Rana contributed to manuscript writing. D. Si, J. Hou, and B. Znosko contributed to manuscript reviewing and modification.

## ACKNOWLEDGEMENTS

We are thankful to all members of the Data Analysis & Intelligent Systems (DAIS) research group and Eugene Cheung for their continuous support and suggestions to the manuscript.

